# Cingulate Cortex Atrophy is Associated with Hearing Loss in Presbycusis with Cochlear Amplifier Dysfunction

**DOI:** 10.1101/574228

**Authors:** Chama Belkhiria, Rodrigo C. Vergara, Simón San Martín, Alexis Leiva, Bruno Marcenaro, Melissa Martínez, Carolina Delgado, Paul H. Delano

**Author notes:** **Correspondence:** Dr. Paul H. Delano, Santos Dumont 999, Hospital Clínico Universidad de Chile, 8380456, Santiago, Chile.

## Abstract

Age-related hearing loss is associated with cognitive decline and has been proposed as a risk factor for dementia. However, the mechanisms that relate hearing loss to cognitive decline remain elusive. Here, we propose that the impairment of the cochlear amplifier mechanism is associated with structural brain changes and cognitive impairment. Ninety-six subjects aged over 65 years old (63 female and 33 male) were evaluated using brain magnetic resonance imaging, neuropsychological and audiological assessments, including distortion product otoacoustic emissions as a measure of the cochlear amplifier function. All the analyses were adjusted by age, gender and education. The group with cochlear amplifier dysfunction showed greater brain atrophy in the cingulate cortex and in the parahippocampus. In addition, the atrophy of the cingulate cortex was associated with cognitive impairment in episodic and working memories and in language and visuoconstructive abilities. We conclude that the neural abnormalities observed in presbycusis subjects with cochlear amplifier dysfunction extend beyond core auditory network and are associated with cognitive decline in multiple domains. These results suggest that a cochlear amplifier dysfunction in presbycusis is an important mechanism relating hearing impairments to brain atrophy in the extended network of effortful hearing.

## Introduction

Age-related hearing loss or presbycusis is characterized by bilateral progressive hearing loss and impaired speech understanding, especially in noisy environments (Gates and Mills, 2005). According to recent epidemiological data from the US more than 50% of people aged over 70 years have presbycusis (Goman and Lin, 2016), while due to population aging, it is expected that by the year 2025 there will be more than one billion people with hearing loss in the world (Sprinzl and Riechelmann, 2010). The increasing number of hearing loss patients is alarming, since epidemiological evidence has shown an association between presbycusis and cognitive decline in elderly people (Lin and Albert, 2014; Thomson et al., 2017). For instance, Lin et al. (2011) have shown that individuals with mild to moderate presbycusis have worse results in executive function and psychomotor processing, while other studies have shown that hearing loss is significantly related to global cognitive decline, which can lead to social isolation and depression (Harrison Bush et al., 2015; Panza et al., 2015). Furthermore, a recent prospective cohort has reported that presbycusis subjects with audiometric hearing thresholds worse than 40 dB are more likely to develop dementia (Deal et al., 2017). In line with this evidence, a Lancet consortium recently proposed a model in which hearing loss is the major potentially preventable risk factor for dementia (Livingston et al., 2017).

Although epidemiological and clinical studies confirm the relationship between hearing loss and cognitive decline, the mechanisms that relate them remain elusive (Lin and Albert, 2014; Wayne and Johnsrude, 2015). The pathological correlate of presbycusis can display different features, including cochlear hair cell loss, stria vascularis atrophy, and auditory-nerve neuron loss (Gates and Mills, 2005). Animal models of presbycusis have shown that in addition to the peripheral auditory damage, aged animals have less GABAergic neurons in the auditory cortex (Martin del Campo et al., 2012). In the same line, several human studies have found brain structural changes in patients with hearing loss, including grey matter volume reduction in the right temporal lobe (Lin et al., 2014a; Peelle and Wingfield, 2016) and correlations between hearing impairment and smaller gray matter volume in the auditory cortex (Eckert et al., 2012; Peelle et al., 2011). However, whether the atrophy of specific brain regions and cognitive domains in presbycusis patients are associated with cochlear receptor cell loss is unknown (Shen et al., 2018).

There are two types of receptor cells in the mammalian cochlea: inner hair cells (IHC) and outer hair cells (OHC). While, IHCs are related to sensory transduction of acoustic vibrations, OHCs are responsible for the cochlear amplifier mechanism, which allows high sensitivity and sharp frequency selectivity to low-intensity sounds (Brownell et al., 1985; Robles and Ruggero, 2001; Liberman et al., 2002). In natural hearing, the cochlear amplifier is fundamental for frequency discrimination and speech perception, and can be assessed through otoacoustic emissions.

Here we hypothesized that a cochlear amplifier dysfunction in presbycusis –an OHC loss–is associated with cognitive decline and structural brain changes in the elderly population. We studied the possible associations between cochlear OHC function (measured by distortion product otoacoustic emissions [DPOAE]), cognitive performance, and brain structure (magnetic resonance imaging, MRI) in a Chilean cohort of elders without dementia (ANDES) and with different levels of hearing loss.

## Materials and methods

All procedures were approved by the Ethics Committee of the Clinical Hospital of the University of Chile, permission number: OAIC752/15. All subjects gave written informed consent in accordance with the Declaration of Helsinki.

### Subjects

For this cross-sectional prospective study, a total of 142 patients aged ≥ 65 years (screened between 2016 and 2018) from Recoleta’s primary health public center located in the city of Santiago (Chile) were evaluated for recruitment. According to the inclusion and exclusion criteria of the Auditory and Dementia study (ANDES), 96 subjects were included for neuropsychological, audiological and brain imaging assessments. Inclusion criteria were the following: patients older than 65 years without dementia, which was evaluated by a mini-mental state examination (MMSE) score ≥ 24 and preserved functionality measured by the Pfeffer activities questionnaire (Quiroga et al., 2004). Exclusion criteria of recruitment in the study were the following: (i) stroke or other neurological disorders; (ii) dementia; (iii) major psychiatric disorders; (iv) other causes of hearing loss different from presbycusis (e.g. conductive hearing loss); (v) patients using hearing aids and (vi) other causes of significant disability.

### Audiological evaluations

All the audiological evaluations were carried out by an experienced audiologist in a double wall soundproof room located in the Otolaryngology Department of the Clinical Hospital of the University of Chile. Air conduction pure tone audiometric (PTA) hearing thresholds were evaluated at 0.125, 0.25, 0.5, 1, 2, 3, 4, 6 and 8 kHz for each subject in both ears using a clinical audiometer (AC40, Interacoustics®). In addition, in order to rule out conductive hearing loss, bone conduction thresholds were measured at 0.25, 0.5, 1, 2, 3 and 4 kHz. All audiometric data presented in the present work refer to air conduction thresholds. The PTA at 0.5, 1, 2 and 4 kHz were calculated for each subject in both ears. Subjects were classified according to their hearing level: normal hearing (≤ 20 dB), mild presbycusis (> 20 dB, and ≤ 35 dB), and moderate presbycusis (>35 dB) based on the PTA of the better hearing ear.

### Distortion product otoacoustic emissions

Otoacoustic emissions are sounds emitted by the OHCs of the inner ear that can be measured through sensitive microphones (Kemp, 1978). DPOAEs are a type of otoacoustic emission, commonly measured in clinical and basic research as a non-invasive tool for assessing the OHC function of the cochlear receptor (Abdala and Visser-Dumont, 2001). In this study, DPOAE (2f1–f2) were measured as a proxy of the cochlear amplifier function, using an ER10C microphone (Etymotic Research^®^), presenting eight pairs of primary tones (f1 and f2, at 65 and 55 dB SPL, f2/f1 ratio of 1.22) in each ear at eight different 2f1–f2 frequencies: 707 Hz, 891 Hz, 1122 Hz, 1414 Hz, 1781 Hz, 2244 Hz, 2828 Hz and 3563 Hz. Figure 1A shows an example of the spectrogram of the microphone signal obtained from the external ear canal of the subjects, in which a 2f1-f2 DPAOE peak can be observed. To consider the presence of a detectable DPOAE, we used an amplitude criterion: the amplitude of a given DPOAE (dB SPL) should be at least 6 dB above noise floor. Using this criterion, we counted the number of detectable DPOAEs per ear, going from 0 to 8: “0” meaning that the subject had no detectable DPOAE in that ear, and consequently that ear was classified as having cochlear amplifier dysfunction (see below), while “8” meaning that the subject had detectable DPOAEs at all tested frequencies in that ear (normal cochlear amplifier function); and “16” meaning that the subject had detectable DPOAE in all tested frequencies in both ears. In contrast with a method that only measures the amplitude of DPOAE, the procedure of counting the number of detectable DPOAE per ear allowed us to evaluate the cochlear amplifier function in the whole sample (n=96), without eliminating subjects with no detectable DPOAEs (and no measurable amplitude in dB SPL), which is common in elderly subjects.

**Figure 1:**
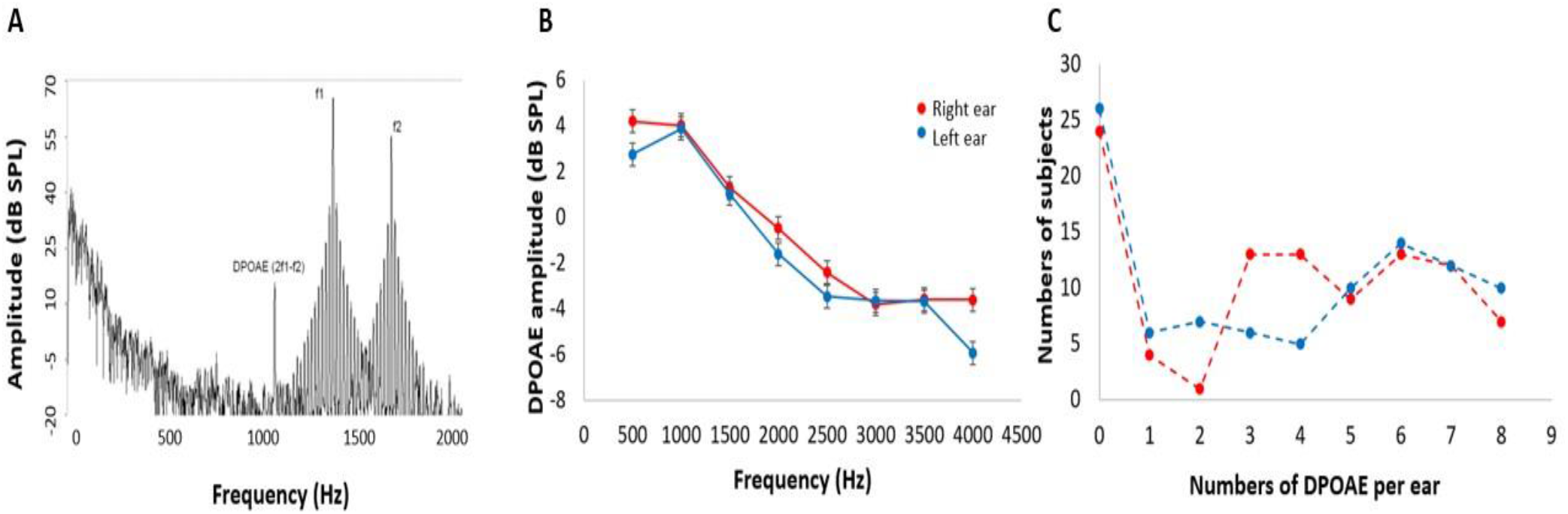
DPOAE measurements. **(A)** Example of the spectrogram of the microphone signal used to obtain the 2f1-f2 DPOAEs in response to f1 and f2 stimuli. DPOAE amplitudes are about 40-50 dB smaller than f1 and f2 primary tones. **(B)** Mean DPOAE amplitudes in response to the eight frequencies for left (blue) and right (red) ears (data shown as mean ± SEM). Notice that low frequency DPOAEs are larger in amplitude than high frequency DPOAEs. **(C)** Frequency histogram of the number of detectable DPOAE per subject for left (blue) and right (red) ears. Note that 24 and 26 subjects had no detectable DPOAEs in right and left ears respectively.

### Cognitive assessment

All subjects and their relatives were evaluated by a neurologist during a structured interview and graded according to their cognitive complaints using the clinical dementia rating scale (Morris, 1993). Cognitive performances were assessed by an experienced psychologist in cognitive tests adapted for the Chilean population (Quiroga et al., 2004), including the Mini-Mental State Examination (MMSE) for global cognition (Folstein et al., 1975), the Frontal Assessment Battery (FAB) for measuring executive function (Dubois et al., 2000), the Trail Making Test A (TMT A) for processing speed (Army Individual Test Battery, 1944), the Boston Nominating Test for Language (Kaplan, 1978), the Rey-Osterrieth Complex Figure Test for Visuospatial Abilities (Rey, 1959), the Backward Digit Span for Verbal Working Memory, and the total recall of the Free and Cued Selective Reminding Test (FCSRT) to explore verbal episodic memory (Grober et al., 1988). Depression was self-rated by the participants using the Geriatric Depression Scale (GDS), which consists of 15 YES/NO answer questions. Higher scores indicate higher levels of depression, with scores over five indicating significant depressive symptoms (Yesavage et al., 1983).

### Magnetic resonance imaging

Neuroimaging data were acquired by a MAGNETOM Skyra 3-Tesla whole-body MRI Scanner (Siemens Healthcare GmbH®, Erlangen, Germany) equipped with a head volume coil. T1-weighted magnetization-prepared rapid gradient echo (T1-MPRAGE) axial images were collected, and parameters were as follows: time repetition (TR) = 2300 ms, time echo (TE) = 232 ms, matrix = 256 × 256, flip angle = 8°, 26 slices, and voxel size = 0.94 × 0.94 × 0.9 mm^3^. T2-weighted turbo spin echo (TSE) (4500 TR ms, 92 TE ms) and fluid attenuated inversion recovery (FLAIR) (8000 TR ms, 94 TE ms, 2500 TI ms) were also collected to inspect structural abnormalities. A total of 440 images were obtained during an acquisition time of 30 minutes per subject.

### Morphometric Analyses

To determine the structural brain changes of controls and presbycusis individuals, we measured the volume and thickness of bilateral cortical regions. FreeSurfer (version 6.0, http://surfer.nmr.mgh.harvard.edu) was used with a single Linux workstation using Centos 6.0 as was suggested by Gronenschild et al. (2012) for T1-weighted images analysis of individual subjects. The FreeSurfer processing involved several stages, as follows: volume registration with the Talairach atlas, bias field correction, initial volumetric labeling, nonlinear alignment to the Talairach space, and final volume labeling. We used the “recon-all” function to generate automatic segmentations of cortical and subcortical regions. This command performs regional segmentation and processes gross regional volume in a conformed space (256×256×256 matrix, with coronal reslicing to 1 mm^3^ voxels). The function “recon-all” creates gross brain volume extents for larger-scale regions (i.e., total number of voxels per region): total grey and white matter, subcortical grey matter, brain mask volume, and estimated total intracranial volume.

Additionally, we measured the cortical thickness in native space using Freesurfer tools. We calculated the cortical thickness of each mesh of vertices by measuring the distance between the point on one surface and the closest conforming point on the opposite surface. Then we measured the average of the two values calculated from each side to the other (Fischl and Dale, 2000). Based on the brain regions that have been previously studied in presbycusis (Lin et al., 2014b; Rigters et al., 2017) our regions of interest were the bilateral frontal superior, temporal inferior, middle and superior gyri, parahippocampus, hippocampus, and amygdala. We also included as regions of interest, cortical areas that have been implicated in the neural networks of degraded speech comprehension: bilateral anterior cingulate cortex (ACC), posterior cingulate cortex (PCC), and precentral and postcentral gyri (Peelle and Wingfield, 2016).

### Statistical analysis

SPSS software version 20.0 was used to perform statistical analysis of demographic data, audiometric measurements, neuropsychological test results and structural MRI data. The ANCOVA method was used to compare cognitive performances and volume and thickness differences between the three tested groups including age, sex, and years of education as covariables. Pearson correlations were performed according to the distribution of data.

We applied general linear models (GLM) to contrast the difference of correlations between the studied groups using PTA as predictor for both volume and thickness. Age, education and sex were adjusted as covariates. The results were corrected for the multiple comparison using a significant alpha value of p = 0.01 (Hagler et al., 2006). Regions with significant differences in GLM models were selected based on the results of multiple corrections, and their corresponding brain regions were represented using the scientific notation p values. The volume and thickness of selected regions were calculated based on the spatially normalized images of each individual in Talairach space (Fischl et al., 2002). In addition, we performed partial correlations including age, sex and education as confound regressors to associate cognitive tests with anterior and posterior cingulate cortex volume and thickness.

## Results

### Demographics characteristics

The mean age of the recruited subjects (n=96, 63 female) was 73.6 ± 5.3 years (mean ± standard deviation [SD]) with an average educational level of 9.5 ± 4.2 years of schooling (mean ± SD). The audiometric hearing thresholds were normal in 32 subjects, while 44 and 20 individuals were considered as mild and moderate presbycusis respectively. None of the 96 subjects used hearing aids at the moment of recruitment. A summary of demographic data, including age, sex, education, hearing level, smoking, and cardiovascular risk factors is presented in Table 1.

**Table 1:** Demographic characteristics of the ANDES cohort.

| Characteristic | Cohort ($n = 96$) |
| --- | --- |
| Age, mean (SD) | 73.62 (5.34) |
| Sex, $n$ (%) | |
| Female | 63 (65.62) |
| Education, mean years (SD) | 9.53 (4.23) |
| Hearing loss, mean dB (SD) | 25.35 (10.91) |
| Hearing loss category, $n$ (%) | |
| Normal, ($<20$ dB) | 32 (33.33) |
| Mild, (21-35 dB) | 44 (45.83) |
| Moderate, ($>35$ dB) | 20 (20.83) |
| DPOAEs, mean number (SD) | 7.45 (5.35) |
| Cochlear status, $n$ (%) | |
| Preserved cochlear amplifier | 64 (66.66) |
| Cochlear amplifier dysfunction | 32 (33.33) |
| Hypertension, $n$ (%) | 39 (40.62) |
| Smoking, $n$ (%) | 18 (18.75) |
| Diabetes, $n$ (%) | 25 (26.04) |
| Hearing aid use, $n$ (%) | 0 (0.0) |

### Cochlear amplifier function

Figure 1B shows the mean amplitudes of the DPOAEs measured at eight different frequencies per ear, while Figure 1C displays a frequency histogram of the number of detectable DPOAE per ear, showing a bimodal distribution with a peak below two DPOAE per ear, and a broad distribution for more than two detectable DPOAE per ear. According to this bimodal distribution of the number of detectable DPOAE, we decided to split data into two groups: (i) better preserved cochlear amplifier function (≥5 DPOAE in both ears) and (ii) cochlear amplifier dysfunction (<5 DPOAE in both ears). Figure 2 shows that the number of detectable DPAOE significantly correlates with age and audiometric hearing thresholds, meaning that a lower number of DPOAE is associated with more aged subjects and worse hearing thresholds.

**Figure 2:**
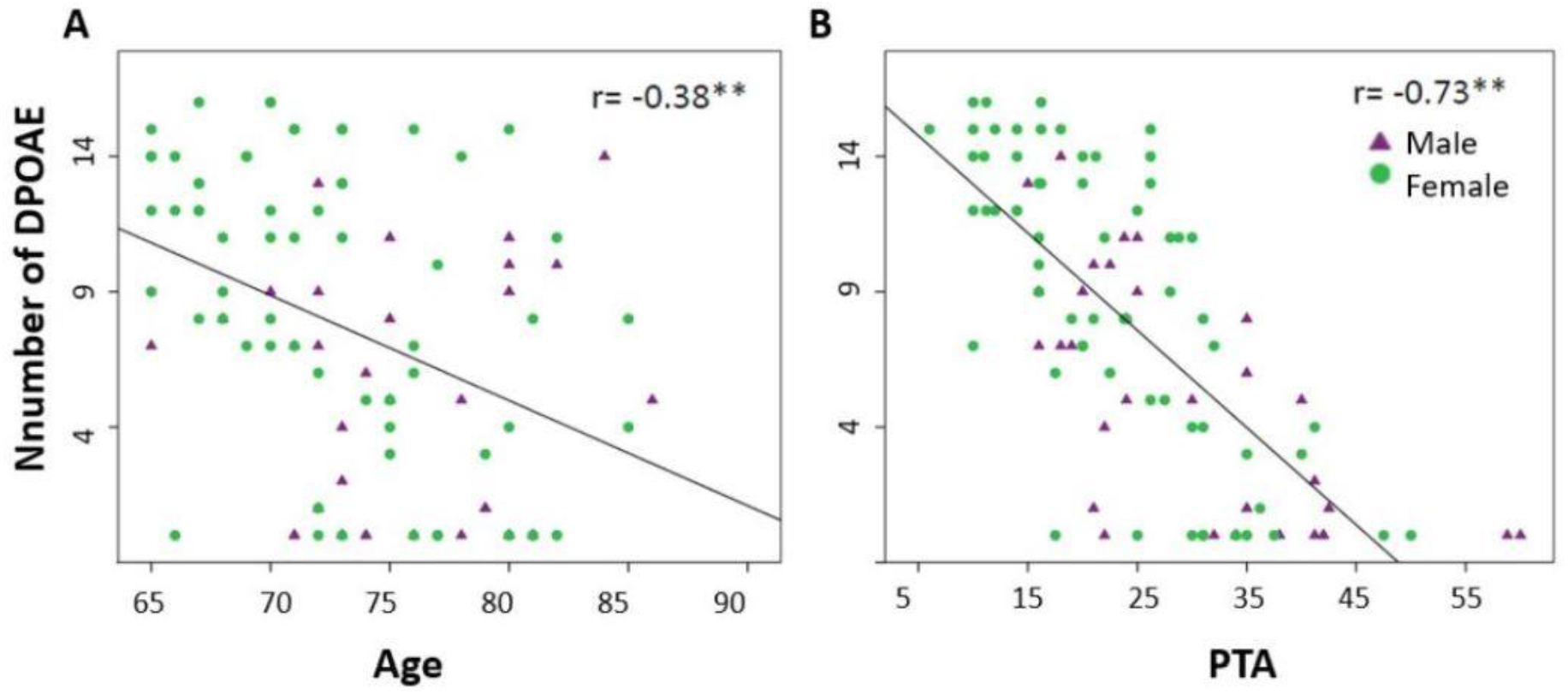
The number of detectable DPOAEs correlates with age and hearing thresholds (PTA). **(A)** A significant negative Spearman correlation was found between age and the number of detectable DPOAE in both ears (**p<0.05, r=−0.38). **(B)** A significant negative Spearman correlation was found between the audiometric thresholds of the better hearing ear (PTA) and the number of detectable DPOAE in both ears (**p<0.05, r=−0.73). Green circles and purple triangles represent female and male subjects, respectively.

Next, in addition to the classification based on the cochlear amplifier status, we used the degree of hearing loss (PTA) to divide subjects into three groups: (i) Controls: normal hearing (≤ 20 dB PTA, better hearing ear) with preserved cochlear amplifier function (≥ 5 detectable DPOAE) (n = 31); (ii) mild and moderate presbycusis (> 20 dB PTA, better hearing ear) with preserved cochlear amplifier (PCF) (n = 33); and (iii) mild and moderate presbycusis (> 20 dB PTA, better hearing ear) with cochlear amplifier dysfunction (<5 detectable DPOAE) (CD) (n = 31). One subject could not be classified within these criteria as he presented normal hearing with cochlear amplifier dysfunction and was excluded for further analyses.

Table 2 shows a summary of the audiological and cognitive performance of the three groups (controls, PCF and CD) controlled by age, gender and years of education. There was a significant difference in age between controls and the group of presbycusis with cochlear amplifier dysfunction, while non-significant differences in age and education were found between the two presbycusis groups (PCF and CD). At the cognitive level, the group with presbycusis and cochlear amplifier dysfunction had worse executive function than the other groups.

**Table 2.**
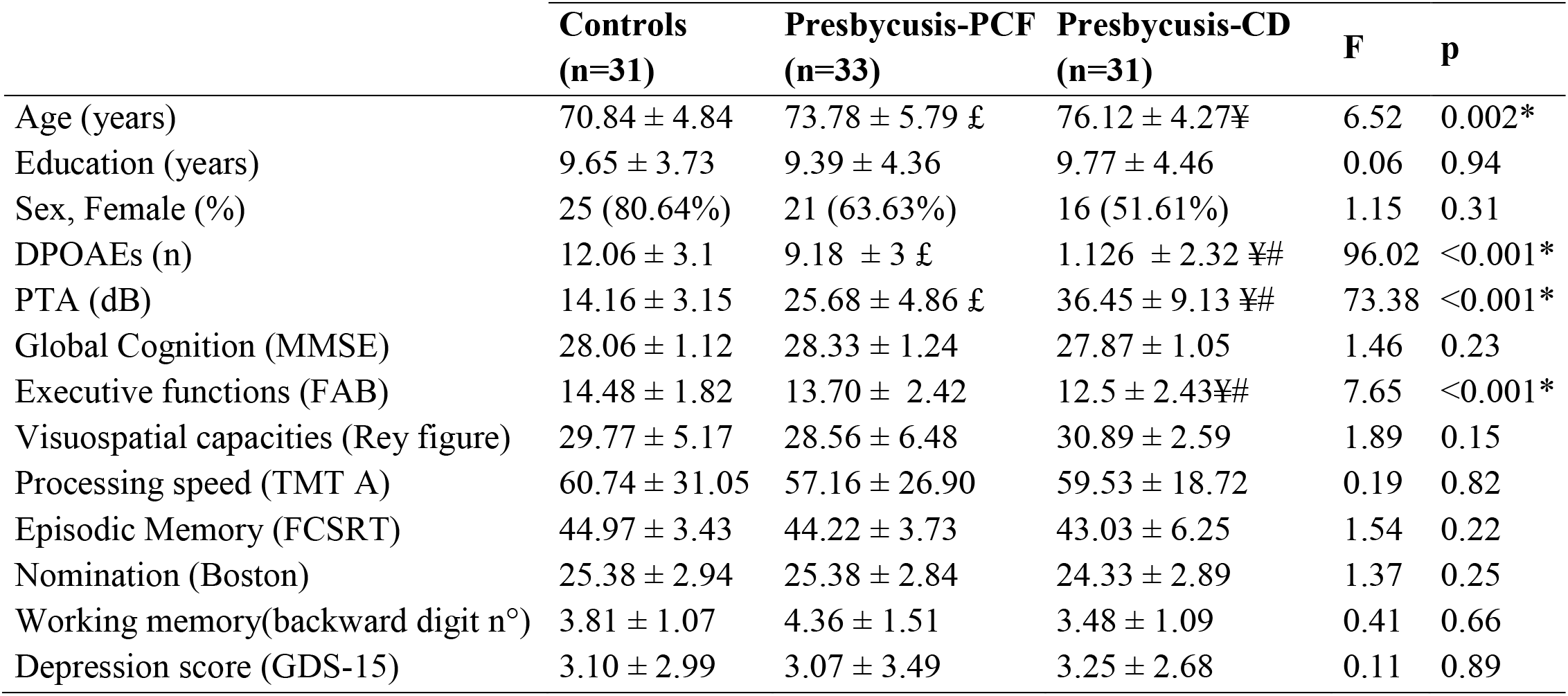
Demographic, audiological and cognitive differences between controls and presbycusis groups with and without cochlear amplifier dysfunction. Data were analyzed using ANCOVA for comparing variables between the three groups corrected by age, sex and education. Mean ± SD values for age, education, sex, DPOAEs, PTA, cognitive tests are shown for controls, presbycusis with preserved cochlear function (PCF) and presbycusis with cochlear amplifier dysfunction (CD) groups. * p < 0.05 for significant ANCOVA. ¥ p < 0.05 significantly different when comparing presbycusis with cochlear amplifier dysfunction vs controls. # p < 0.05 significantly different when comparing presbycusis with CD vs presbycusis without CD. £ p < 0.05 significantly different when comparing presbycusis without CD vs controls.

### Structural brain changes

Our next step was to evaluate whether structural brain changes were associated with cochlear amplifier dysfunction. Hence, we compared the MRI volumes and thickness of cortical and limbic brain regions in the three groups (controls, PCF and CD) using ANCOVA models adjusted by age, sex and education. Structural volumes were normalized by the total estimated intracranial volume. The ANCOVA models for brain volumes yielded significant differences between the three groups in the precentral (F(2,89)=3.92, p=0.02) and postcentral gyri (F(2,89)=3.92, p=0.02), parahippocampus (F(2,89)=4.55, p=0.01) and amygdala (F(2,89)=3.31, p=0.04). Tukey post hoc tests showed significant differences between the CD and controls in the volume of precentral and postcentral gyri and parahippocampus. The only significant difference in brain volume between PCF and CD groups was obtained from parahippocampus (Table 3).

**Table 3.** Brain volume and thickness differences between the controls and presbycusis groups with and without cochlear amplifier dysfunction. Data were analyzed using ANCOVA for comparing variables between the three groups corrected by age, sex and education. Mean ± SD values for the sum of bilateral brain region thickness and volumes normalized by total estimated intracranial volume. * p < 0.05 for significant ANCOVA. ¥ p < 0.05 significantly different when comparing presbycusis with CD vs controls. # p < 0.05 significantly different when comparing presbycusis with CD vs presbycusis without CD.

|  | Controls<br>n=31 | PCF<br>n=33 | CD<br>N=31 | F | p |
| --- | --- | --- | --- | --- | --- |
| **Brain volume regions (cm<sup>3</sup>)** |  |  |  |  |  |
| Anterior cingulate | 2.87 ± 0.4 | 2.74 ± 0.53 | 2.69 ± 0.38 | 1.2 | 0.3 |
| Posterior cingulate | 4.03 ± 0.53 | 3.98 ± 0.59 | 3.94 ± 0.47 | 0.23 | 0.79 |
| Precentral gyrus | 8.01 ± 1.1 | 7.6 ± 1.0 | 7.2 ± 1.0 ¥ | 3.92 | 0.02* |
| Postcentral gyrus | 5.1 ± 0.1 | 4.9 ± 0.7 | 4.5 ± 0.7 ¥ | 3.92 | 0.02* |
| Parahippocampus | 2.9 ± 0.5 | 2.9 ± 0.4 | 2.6 ± 0.4 ¥ # | 4.55 | 0.01* |
| Frontal superior | 13.7 ± 2.2 | 13.3 ± 1.7 | 13.1 ± 1.9 | 0.66 | 0.51 |
| Temporal inferior | 8.8 ± 1.6 | 8.8 ± 0.8 | 8.6 ± 1.9 | 0.25 | 0.77 |
| Temporal middle | 14.4 ± 2.5 | 14.3 ± 1.8 | 13.7 ± 2.5 | 0.89 | 0.41 |
| Temporal superior | 20.4 ± 0.2 | 20.5 ± 0.2 | 19.8 ± 0.2 | 0.59 | 0.55 |
| Amygdala | 2.2 ± 0.3 | 2.2 ± 0.3 | 2.01 ± 0.4 | 3.31 | 0.04* |
| Hippocampus | 5.5 ± 0.8 | 5.3 ± 0.7 | 5.01 ± 0.8 | 2.41 | 0.09 |
| **Cortical thickness (mm)** |  |  |  |  |  |
| Anterior cingulate | 4.81 ± 0.26 | 4.78 ± 0.33 | 4.48 ± 0.32 ¥# | 3.04 | 0.04* |
| Posterior cingulate | 4.99 ± 0.18 | 4.97 ± 0.25 | 4.6 ± 0.28 ¥# | 4.17 | 0.01* |
| Precentral gyrus | 4.97 ± 0.24 | 4.9 ± 0.24 | 4.79 ± 0.29 ¥ | 4.00 | 0.02* |
| Postcentral gyrus | 4.08 ± 0.21 | 3.99 ± 0.18 | 3.91 ± 0.21 ¥ | 6.43 | 0.002* |
| Parahippocampus | 5.36 ± 0.58 | 5.33 ± 0.48 | 5.06 ± 0.57 | 3.30 | 0.04* |
| Frontal superior | 4.91 ± 0.18 | 4.91 ± 0.22 | 4.86 ± 0.25 | 0.54 | 0.58 |
| Temporal inferior | 5.51 ± 0.19 | 5.52 ± 0.26 | 5.48 ± 0.26 | 0.23 | 0.79 |
| Temporal middle | 5.32 ± 0.17 | 5.31 ± 0.21 | 5.22 ± 0.22 | 2.07 | 0.13 |
| Temporal superior | 5.33 ± 0.25 | 5.30 ± 0.25 | 5.21 ± 0.3 | 2.08 | 0.13 |

Regarding cortical thickness, we found significant differences in the ANCOVA models for anterior cingulate cortex (F(2,89)=3.04, p=0.04), posterior cingulate cortex F(2,89)=4.17, p=0.01), precentral gyrus (F(2,89)=4.00, p=0.02), postcentral gyrus (F(2,89)=6.43, p=0.002) and parahippocampus (F(2,89)=3.3, p=0.04). Tukey posthoc tests showed that the thickness of precentral and postcentral gyri in the CD group were significantly reduced compared to that of the control group, while the thickness of ACC and PCC in the CD group were thinner than those of the PCF group (Table 3).

Next, in order to isolate the effect of hearing levels and cochlear amplifier dysfunction on brain structures, we ran GLM analyses. The GLM contrasted the difference of associations between PCF and CD groups using PTA as predictor for both volume and thickness. This method can effectively select voxels that correlate with PTA when contrasting PCF and CD groups. The whole brain analyses of volume and thickness related to cochlear amplifier dysfunction revealed significant correlations between PTA and different brain regions (corrected by age, education and sex, and using a level of significance = p <0.01) shown in red and blue colors in Figure 3. These associations were significant for large clusters of voxels in bilateral ACC and PCC. Cochlear amplifier dysfunction was also associated with significant negative correlations between PTA and volume and between PTA and thickness in scattered regions of the right superior temporal, right postcentral and left precentral areas. We also found small clusters resulting from positive correlation between PTA and volume in the right superior and middle frontal areas in the CD group (Figure 3A).

**Figure 3:**
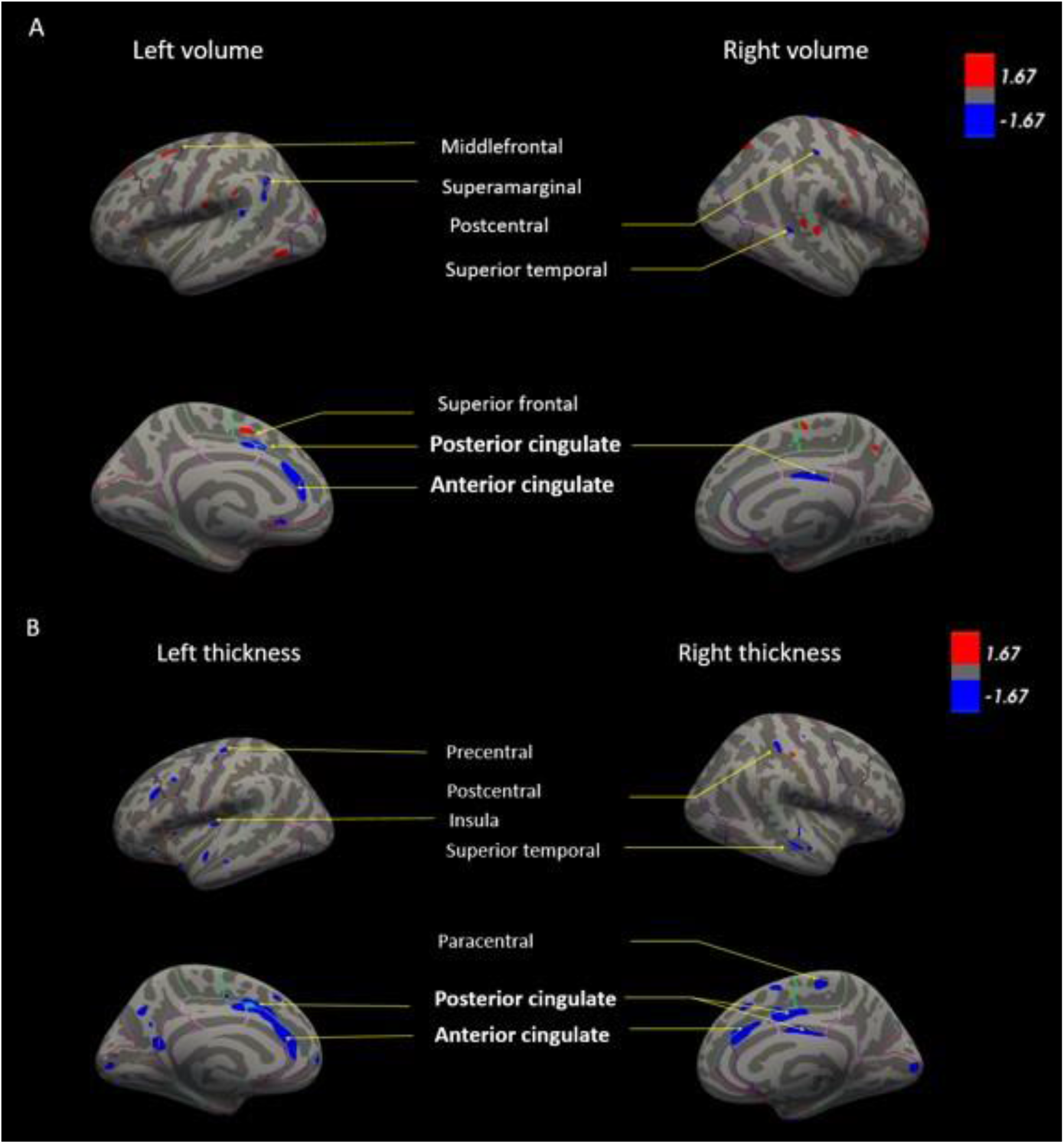
GLM models for brain volume (A) and cortical thickness (B) correlated with PTA in presbycusis patients. The GLM contrasts the difference of correlations between PCF and CD groups using PTA as predictor for both volume and thickness. Age, education and sex were adjusted as covariates. The color blue shows regions in which individuals with cochlear dysfunction had a higher rate of gray matter decrease compared to those with normal cochlear (p< 0.01). The color red shows regions in which individuals with cochlear impairment had a higher rate of gray matter increase compared to those with normal cochlear (p <0.01).

Figure 4 shows examples of the correlations between PTA and bilateral ACC thickness in CD and PCF groups, illustrating that PTA was negatively correlated with the thickness of bilateral ACC gyri only in the CD group.

**Figure 4:**
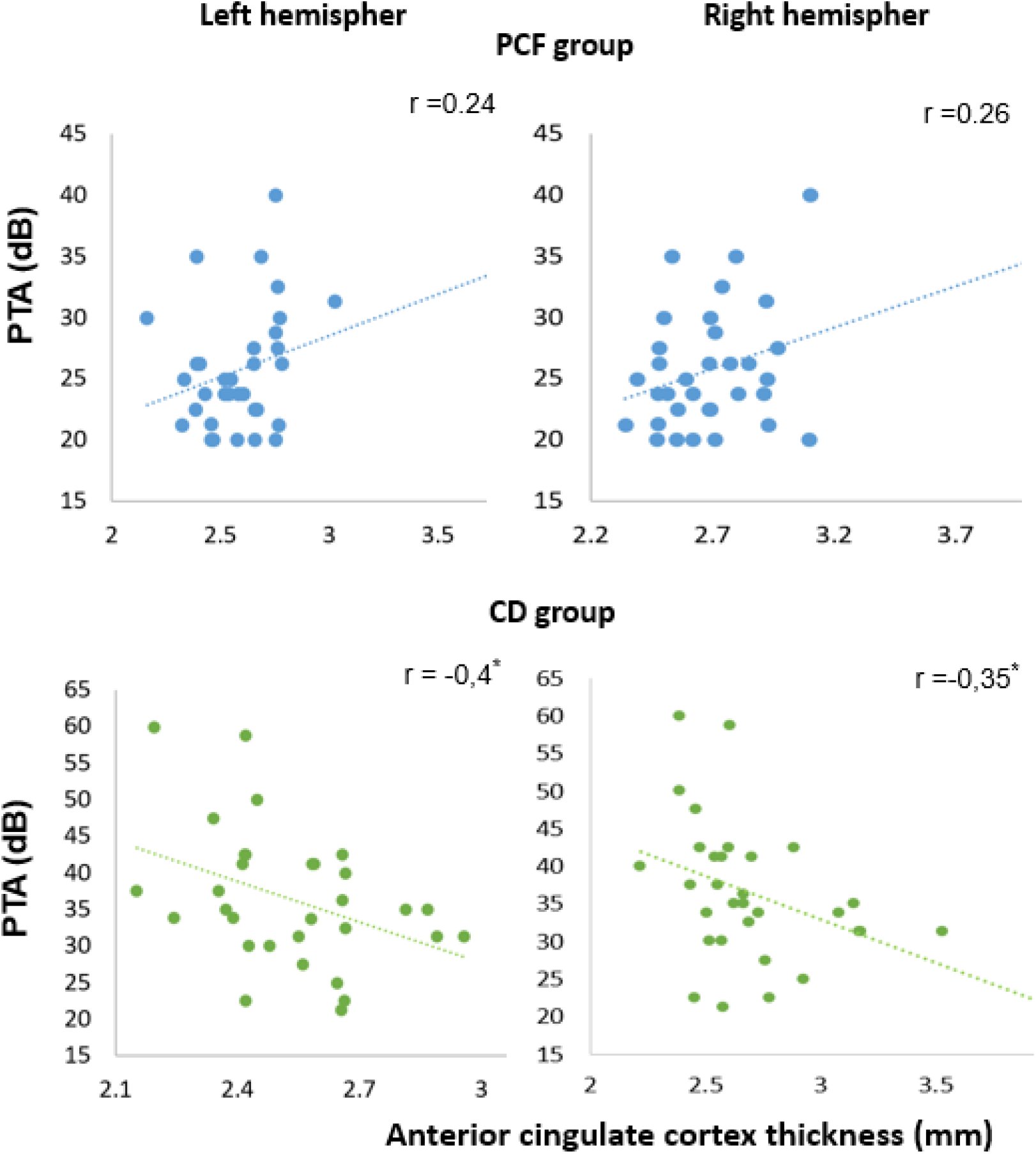
Significant negative correlations between PTA and ACC thickness in presbycusis patients with cochlear amplifier dysfunction. Pearson correlations between PTA and anterior cingulate thickness for PCF (blue) and CD (green) groups.

### Cognitive performance, depression scores and cingulate cortices volume and thickness

Pearson partial correlation analyses were performed between the bilateral volume and thickness of ACC and PCC and cognitive performance in PCF and CD groups. Age, sex and education were controlled as covariates (Table 4). Correlations between cognitive tests and structural measures of ACC and PCC were different between CD and PCF groups. In CD patients, there were significant correlations between episodic memory (total recall in the FCSRT), nomination, verbal working memory tests and volume of the left ACC. On the other hand, processing speed and verbal working memory correlated with the volume of the left PCC, while visuospatial abilities with the right PCC. These results indicate that, in the group with cochlear amplifier dysfunction, the volume of cingulate cortices and cognition performance decreased together. In addition, there was a negative correlation between depression scores and ACC volume in the CD group, indicating that higher depressive symptoms were associated with a decreased ACC volume.

**Table 4.** Pearson partial correlation corrected by age, sex, and education between cognitive tests and volume and thickness of anterior and posterior cingulate cortex for PCF and CD groups (*p<0.05).

| Cognitive tests | ACC volume |  |  |  | ACC thickness |  |  |  |
| --- | --- | --- | --- | --- | --- | --- | --- | --- |
|  | PCF |  | CD |  | PCF |  | CD |  |
|  | r-Left | r-Right | r-Left | r-Right | r-Left | r-Right | r-Left | r-Right |
| Cognitive functions (MMSE) | 0.14 | 0.46* | -0.19 | 0.00 | 0.22 | 0.17 | 0.03 | -0.13 |
| Executive functions (FAB) | 0.29 | 0.07 | -0.01 | 0.21 | 0.14 | 0.26 | 0.62 | 0.13 |
| Visuospatial capacities (Rey figure) | 0.05 | 0.41* | 0.14 | 0.30 | 0.01 | 0.28 | 0.15 | 0.49* |
| Processing speed (TMT A) | -0.27 | 0.01 | -0.37 | -0.04 | -0.22 | -0.17 | -0.34* | -0.26 |
| Episodic Memory (FCSRT) | -0.20 | 0.19 | 0.44* | 0.31 | 0.01 | 0.08 | 0.34 | 0.10 |
| Nomination (Boston) | 0.21 | 0.03 | 0.47* | 0.29 | 0.03 | 0.07 | 0.48* | 0.41* |
| Working memory (backward digit n°) | 0.10 | -0.14 | 0.64* | 0.40 | -0.06 | -0.42 | 0.18 | 0.07 |
| Depression score (GDS-15) | 0.32 | -0.20 | -0.53* | -0.01 | -0.17 | 0.36 | 0.05 | 0.00 |
| Cognitive tests | PCC volume |  |  |  | PCC thickness |  |  |  |
|  | PCF |  | CD |  | PCF |  | CD |  |
|  | r-Left | r-Right | r-Left | r-Right | r-Left | r-Right | r-Left | r-Right |
| Cognitive functions (MMSE) | 0.05 | 0.40 | -0.10 | 0.00 | 0.14 | 0.14 | -0.01 | -0.19 |
| Executive functions (FAB) | -0.12 | -0.13 | -0.01 | 0.02 | -0.03 | 0.04 | -0.11 | -0.24 |
| Visuospatial capacities (Rey figure) | 0.32 | 0.27 | 0.35 | 0.54* | 0.00 | 0.02 | 0.35 | 0.45* |
| Processing speed (TMT A) | -0.40* | -0.21 | -0.41* | -0.31 | 0.02 | -0.01 | -0.20 | -0.11 |
| Episodic Memory (FCSRT) | -0.26 | 0.05 | 0.28 | 0.22 | 0.05 | 0.11 | 0.08 | -0.18 |
| Nomination (Boston) | 0.19 | 0.11 | 0.23 | 0.31 | -0.16 | 0.03 | 0.28 | 0.20 |
| Working memory (backward digit n°) | 0.10 | 0.05 | 0.45* | 0.18 | 0.12 | 0.40 | -0.35 | 0.00 |
| Depression score (GDS-15) | 0.17 | 0.00 | -0.12 | 0.04 | 0.27 | 0.35 | 0.25 | -0.09 |

Regarding cortical thickness, we found significant correlations between visuospatial capacities and right ACC, processing speed and left ACC, and nomination with bilateral ACC in the CD group. The visuospatial capacities test also correlated significantly with the thickness of the right PCC in the CD group. Taken together, these results indicate that the more severe the cognitive impairment, the thinner the cortex is in the anterior and posterior cingulate area.

## Discussion

In this combined approach, we found that structural brain changes in presbycusis subjects with cochlear amplifier dysfunction extend beyond central auditory pathways, including the cingulo-opercular network which is fundamental for effortful hearing. In addition, these changes were associated with impairments in verbal and non-verbal cognitive domains and depressive symptoms.

### Cochlear amplifier dysfunction and hearing loss

A reduced number of detectable DPOAEs correlated with age, confirming that cochlear damage increases with aging (Abdala et al., 2018; Abdala and Dhar, 2012). In addition, the number of DPOAE and hearing levels were also correlated, which could be explained by the fact that cochlear aging is one of the main causes of presbycusis (Gates and Mills, 2005). However, it is important to highlight that both measures (DPOAE and PTA) are not identical and several differences can be described. DPOAE are objective measures that can be obtained with a microphone positioned in the external ear canal of anesthetized or unconscious subjects, while audiometric thresholds need the voluntary response of individuals. DPOAEs specifically reflect the function of the cochlear amplifier located in the outer hair cells of the cochlear receptor (Robles et al., 1991), while audiometric thresholds depend on the normal function of the complete auditory pathway.

In our work, according to the bimodal distribution of DPOAE presence, we divided data of presbycusis patients into two groups with different levels of cochlear amplifier function. As the cochlear amplifier is essential for auditory sensitivity, frequency and speech discrimination (Robles and Ruggero, 2001), we hypothesized that the group of patients with loss of OHCs would have more severe neural and cognitive consequences of hearing loss. Indeed, our results showed that the CD group had worse executive functions in comparison with the group with PCF, even after adjusting by age, education, and sex. This finding is in agreement with previous reports showing that executive functions are associated with hearing impairment (Lin et al., 2011), but we extended this association showing that executive functions are more impaired in the group with cochlear amplifier dysfunction.

### Cochlear amplifier dysfunction and brain structure

At the brain structural level, we compared the volume and cortical thickness of regions of interest that were previously reported to be related to hearing loss, but we also included areas beyond the temporal lobe, that have been involved in effortful hearing (Peelle 2016). Our results confirm that hearing impairment was associated to right superior temporal volume decrease (Lin et al., 2014a; Rigters et al., 2017), but also to right superior temporal thickness decrease in the CD group. Notably, in the group of patients with CD, hearing loss produced larger effects in the network of effortful hearing, including the ACC, PCC, precentral and postcentral cortices, and the parahippocampus (Figure 3). While these findings are in agreement with a study reporting volume decrease in the right anterior cingulate cortex in a group of middle-aged hearing loss patients (Husain et al., 2011), our results stress the importance of cochlear amplifier function in the neural structures related to the effortful hearing network in aged people.

Previous studies using functional MRI in presbycusis patients have demonstrated activation of non-auditory cortical regions related to the extended cortical network that supports spoken language processing (Peelle et al., 2011; Sharma and Glick, 2016). The cingulo-opercular cortex is one of the principal components of this network (Peelle and Wingfield, 2016). Increased activation of this area is related to better speech comprehension in normal hearing subjects (Peelle, 2018), and in mild hearing loss patients during passive listening (Campbell and Sharma, 2013).

Our work shows that the atrophy of the ACC is associated with cognitive and emotional deficits in presbycusis patients with cochlear amplifier dysfunction. The ACC is a key structure of cognitive and emotional variables necessary for decision making and social behavior that has also been implicated in pain, negative affect, and autonomic networks (Fornito et al., 2004; Shackman et al., 2011). Interestingly, we found that in the CD group, the reduction of the ACC volume was correlated not only with impairments in tests related to language processing (episodic and verbal working memories and nomination), but also with non-language functions such as visuospatial abilities and processing speed (Table 4). The decrease in performance in non-auditory cognitive domains could be explained by an increased listening effort in the group of CD patients, which needs the recruitment of non-auditory brain areas, decreasing resources for other cognitive functions (Peelle, 2018).

Another possible mechanism to explain the relation between episodic memory and semantic impairment with ACC atrophy in presbycusis could be a functional disconnection between the temporal lobe and the cingulate region (Warrington and Weiskrantz, 1982), as the functional connectivity between the cingulate and temporal areas has been correlated with memory performance in patients with mild cognitive impairment. Furthermore, the cingulate region and the parahippocampus gyrus are key components of the default mode network (Greicius et al., 2003; Raichle, 2015) which were significantly atrophied in the group of presbycusis with cochlear amplifier dysfunction. Taken together, our findings suggest that the relations between the ACC atrophy and episodic memory impairment could reflect an impairment of the Papez circuit in presbycusis patients with cochlear amplifier dysfunction (Carlesimo et al., 2011; Tsivilis et al., 2008).

At the emotional level, several functional neuroimaging studies have found hypoperfusion of the ACC cortex in subjects with major depression (Fitzgerald et al., 2008). Accordingly, we found that the volume of the left ACC was correlated with depressive symptoms in the group with cochlear amplifier dysfunction. However, it is important to highlight that our study only included subjects with mild to moderate depressive symptoms that could be interpreted as “negative affect or sadness” (Mayberg et al., 1999).

In our study the volume and thickness of the right PCC was also significantly correlated with visuospatial abilities in the CD group. This could be explained by role of the right PCC and the right hippocampus in the cortical network associated with spatial abilities, including spatial navigation and egocentric orientation (Burles et al., 2018; Hirono et al., 1998), suggesting that presbycusis could contribute to these spatial deficits.

In addition to the atrophy of the limbic areas in the CD group, we found changes in sensorimotor brain regions. The volume and thickness of the precentral and postcentral gyri were significantly reduced in the group with CD in comparison with the two other groups (Figure 3). These results could be explained by aging, as these regions are known to be affected in elderly people with gait disorders. Another possibility is the recruitment of precentral and postcentral gyri for speech perception in situations with degraded acoustic signal as part of the extended network of language processing (Peelle et al., 2010; Peelle and Wingfield, 2016).

### Why is cochlear amplifier dysfunction associated with brain atrophy?

One possible explanation is that the loss of OHC could be a general indicator of accelerated cell death in the nervous system of elderly people (Shen et al., 2018). In this sense, we can speculate that the absence of DPAOE in mild presbycusis patients could be an early risk factor for detecting accelerated neurodegeneration that could lead to cingulate cortex atrophy and executive dysfunction.

Another speculative mechanism is that the constant activation of the effortful listening network could produce an accelerated neurodegeneration of cingulate neurons caused by glutamate excitotoxicity (Adank, 2012; Eckert et al., 2009; Vaden et al., 2015). This mechanism has been proposed in other neuropsychiatric diseases with ACC atrophy including fibromyalgia (Luerding et al., 2008; Cagnie et al., 2014) and psychotic depression (Bijanki et al., 2014; Zhang et al., 2017). Moreover, the constant activation of the cingulate cortices and other regions of the extended language network in presbycusis patients with cochlear amplifier dysfunction could act as a “secondary injury” that triggers neurodegeneration in a susceptible brain (Sperling et al., 2011).

### Limitations of the study

In our study, we have a relatively reduced number of presbycusis patients with moderate or severe hearing loss. This is partially due to the inclusion criterion that requires subjects to have never used hearing aids before the moment of recruitment. This criterion was planned to eliminate possible neural compensatory plasticity associated to the use of hearing devices, but it hindered the recruitment of moderate to severe presbycusis patients (as most of them had used hearing aids previously). Consequently the majority of the study’s subjects have only mild to moderate presbycusis. Another limitation of the present work is that DPOAE loss, aging and brain changes could be associated with other peripheral and central auditory impairments that might affect cognition. These factors include inner hair cell damage, auditory-nerve fiber loss and central auditory processing abnormalities, which could also be associated with DPOAE loss. Finally, the three groups are not perfectly matched by age and PTA. We control this limitation using these and other variables as covariates when needed. This control has its limitations, however, given our conditions, main conclusions should not be affected. Therefore our results are promising, but more research using DPOAEs as predictor for cognitive and neural decline is needed to evaluate the extent of this relation.

## Conclusion

In non-demented patients with mild to moderate presbycusis, cochlear amplifier dysfunction was associated with brain atrophy and cognitive decline. We found structural changes in non-auditory brain areas related to the extended language processing network: the anterior and posterior cingulate cortices, sensorimotor areas and the parahippocampus. The dysfunction of this network is related to impairments in cognitive domains beyond auditory processing.

## Acknowledgements

We thank Dr. Luis Robles for reading a previous version of this manuscript.

